# Gradual labeling with fluorogenic probes for MINFLUX nanoscopy of densely packed structures

**DOI:** 10.1101/2024.05.08.593253

**Authors:** Longfang Yao, Dongjuan Si, Liwen Chen, Shu Li, Jiaxin Guan, Jiong Ma, Qiming Zhang, Jing Wang, Lu Wang, Min Gu

## Abstract

Single-molecule localization in densely packed structures represents a formidable challenge in nanoscopy. Here we develop a gradual labeling method with fluorogenic probes for MINFLUX nanoscopy (GLF-MINFLUX), with which individual proteins in packed structures can be sequential illuminated and quantitatively localized at the nanoscale solely by adjusting probe concentration. With a 1.7-fold localization precision improvement and 2.2-fold faster acquisition, GLF-MINFLUX enables dual-channel nanoscale imaging of densely neuronal microtubules and microfilaments, and three-dimensional quantitative mapping of individual TOM20 proteins in mitochondrial clusters.

## Main

Fluorescence nanoscopy has revolutionized cellular imaging by providing nanoscale spatial details of individual molecules. However, precisely imaging and quantifying proteins in densely packed structures is still an unmet goal due to limitations in localization precision (σ) and “on/off” switching kinetics of fluorophores during single-molecule separation in nanoscopy. MINimal fluorescence photon FLUXes (MINFLUX) nanoscopy emerges as a powerful solution for single molecule separation, enabling sub-1 nm precision in the focal plane and about 2 nm in three dimensions^1–4^. Coupled with “on/off” switchable organic dyes such as Alexa Fluor 647, MINFLUX has enabled the effective imaging and elucidation of cellular structures with low protein density, such as nuclear pores^3, 5^.

However, densely packed structures like microtubules within neuronal axons present significant challenges for MINFLUX nanoscopy. In MINFLUX imaging, only one emitter should be present within the donut-shaped excitation beam to enable precise localization (Fig. 1a). Conventional fluorophores like Alexa Fluor 647 commonly exhibit an on/off duty cycle exceeding 0.0005, denoting one fluorescent event per 2000 molecules^6^. Nevertheless, within a circular region of approximately 648 nm in diameter within the excitation beam, the presence of over 8000 probes in axonal microtubules easily engenders multiple emitters, resulting in frequent localization failures (Supplementary Note 1). This compromises crucial information on targeted proteins and significantly prolongs the acquisition time. Developing novel dyes with lower on/off duty cycles or photoactivable probes requires intricate synthesis and optimization^5, 7–10^. An additional challenge arises from the unpredictable and uncontrollable “on/off” switching kinetics of labeled fluorophores. In densely packed structures, targets fall within the localization precision range, resulting in the high overlap of clusters of localization events. Meanwhile, the employment of blinking dyes like Alexa Fluor 647 frequently induces cycles of “on/off” switching, rendering the distinction between overlapped targets unattainable (Fig. 1b). Thus, a method to effectively regulate the number of illuminated targets is imperative for MINFLUX nanoscopy of densely packed structures.

**Fig. 1.**
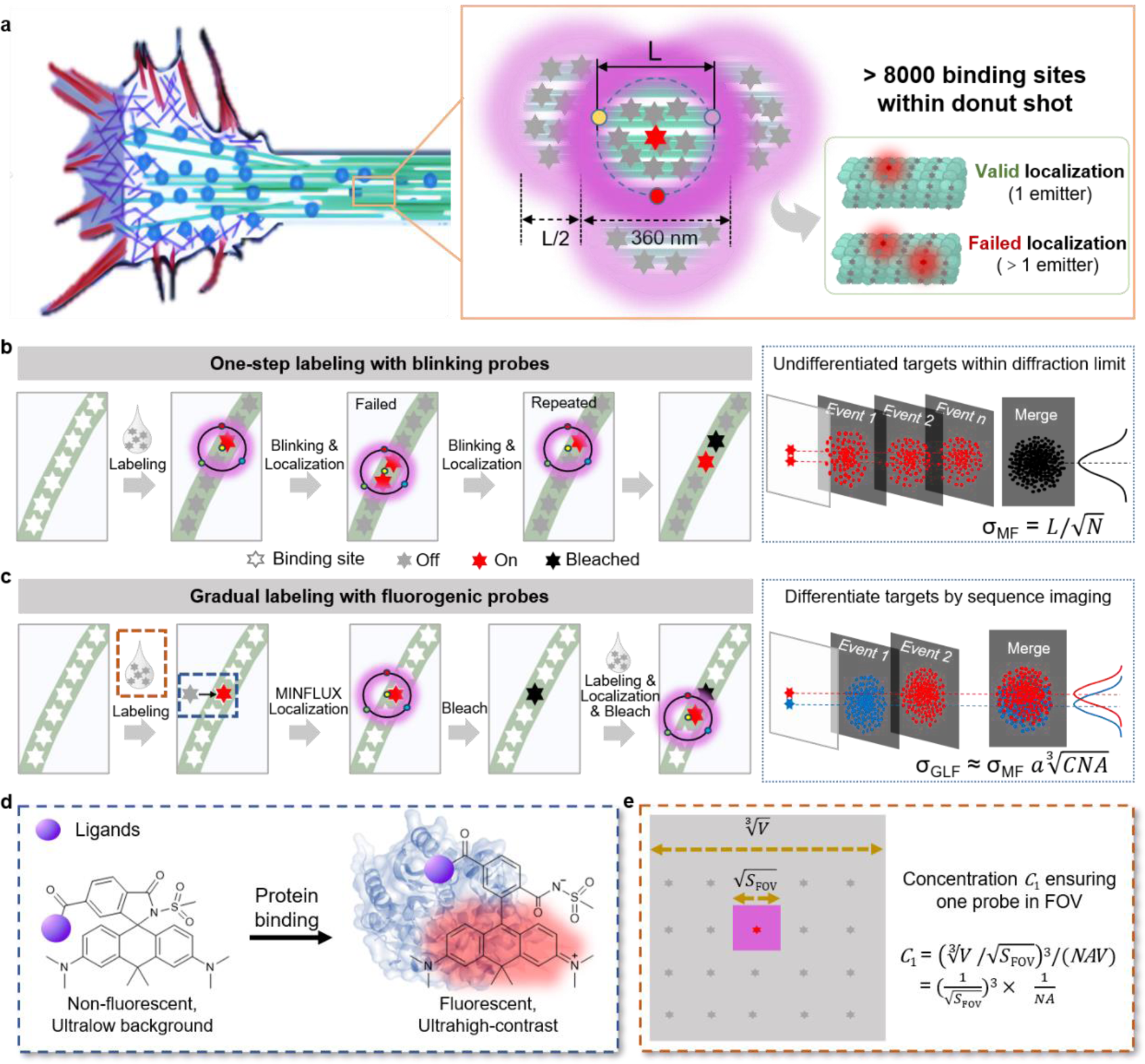
| Schematic diagram illustrating the working principle of GLF-MINLUX. **a2** The scheme indicating the challenge in MINFLUX nanoscopy of densely packed structures like neuronal microtubules. **b**, Working principle of MINLFUX nanoscopy relying on one-step labeling with blinking probes like Alexa Fluor 647 and its challenges in localization failure and differentiation of targets within the localization precision range. **c**, Working mechanism of GLF-MINFLUX. Each valid event encompassing labeling, localization, and bleaching represents a single target protein. The targets within the localization precision range can be segregated and precisely positioned by the iterative labeling and localization steps. **d**, Off-on switching process of **MaP618m** probes upon binding to target protein. **e**, Schematic representation detailing the computational methodology for determining *C*_1_. Area of the imaging field of view (*S_FOV_*), Avogadro’s constant (*NA*), and the volume (*V*).

To alleviate constraints in the “on/off” switching kinetics of organic fluorophores, DNA-based Point Accumulation for Imaging in Nanoscale Topography (DNA-PAINT) has been employed^4^. Importantly, DNA-PAINT combination with resolution through sequential imaging (RESI) allows breakthrough resolution down to the Ångström scale^11^. However, mitigating background noise from diffusing imager strands requires significantly reducing the pinhole diameter, consequently prolonging the imaging duration^4^. The need for abundant orthogonal DNA sequences and meticulous washing and labeling procedures in RESI presents obstacles to nanoscopy in densely packed structures^11^.

In this work, we aim to effectively regulate the number of illuminated targets by introducing **G**radual **L**abeling with **F**luorogenic probes for MINFLUX nanoscopy (GLF-MINFLUX) (Fig. 1c). This method relies on the protein-induced “off/on” switching of **MaP** dye and the resulting high-contrast labeling(Fig. 1d, Extended Data Fig. 1 and Supplementary Table 1, Note 2)^12^. **MaP** probes, in their unbound state, exist as a nonfluorescent spirocyclic compound, transitioning into a fluorescent zwitterion upon binding to target proteins. This protein-induced off-on switching yields exceptional imaging contrast for single-molecule localization without washing out excess unbound probes, or specific buffer systems, or dedicated activation lasers. Thus, by adjusting solely the probe concentration (*C*), we can effectively regulate the number of illuminated targets, ensuring the presence of the single emitter within the field of view in MINFLUX nanoscopy. Importantly, the gradual labeling process enables the utilization of stronger laser power, resulting in a substantially higher photon count and consequently, improved localization precision^13^. Moreover, the employment of a strong excitation laser in GLF-MINFLUX ensures that each valid event, encompassing labeling, localization, and bleaching, represents a single target (Fig. 1c). Consequently, the targets within the localization precision range can be segregated and precisely positioned by the iterative labeling and localization steps within GLF-MINFLUX. Additionally, such gradual labeling and sequential illumination processes also circumvent interference from the coactivation and quenching among adjacent fluorophores. Notably, in comparison to the localization precision of MINFLUX (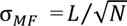),^13^ GLF-MINFLUX possesses a theoretically elevated localization precision 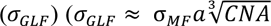, wherein *C* denotes the probe concentration) (Fig. 1c and Supplementary Note 3).

We initially introduced a computational formula 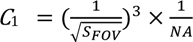 for ascertaining the probe concentration (*C*_1_) (Fig. 1e and Supplementary Note 4), at which the single emitter within the laser beam is ensured. In practical imaging scenarios, a slight elevation in concentration relative to *C*_1_ was implemented and refresh probe solution every 13-20 minutes to increase the labeling rate and minimize imaging time (**Extended Data Fig. and** Supplementary Note 5). To showcase the applicability of GLF-MINFLUX, we stained tubulin-HaloTag-expressing BS-C-1 cells with fluorogenic probe **MaP618m-Halo**^12^, alongside the commonly employed Alexa Fluor 647 as a control (Fig. 2a,b). Conventional one-step labeling using Alexa Fluor 647 yields 9.1 localizations per event and a photon count of 5084 with a reported laser power of 49 µW (Extended Data Fig. 3,4 and Supplementary Note 6) ^3^. Notably, GLF-MINFLUX with **MaP618m-Halo** permits a stronger laser power of 293 µW, resulting in a photon count of 14103 while maintaining 11.9 localizations per localization cloud (Extended Data Fig. 3,4 and Supplementary Note 6). Such a remarkable 2.7-fold increase in photon count in GLF-MINFLUX leads to a 2.6 nm localization precision, which is 1.7-fold higher than Alexa Fluor 647 (Fig. 2c). Importantly, GLF-MINFLUX enables nanoscale imaging of consecutive microtubules within 1.5 hours, whereas only discontinuous microtubules were obtained using one-step labeling with Alexa Fluor 647, even after approximately twice the acquisition time under the same size imaging field of view (Fig. 2a,b,d, Extended Data Fig. 5, Movie 1-2, and Supplementary Note 6). The superior performance from GLF-MINFLUX compared to conventional one-step labeling with Alexa Fluor 647 is further demonstrated by the 2.2-fold higher acquisition rate (Supplementary Note 6), the 1.2-fold higher center-frequency ratio (CFR) (Fig. 2e), a 1.8-fold reduction in the time between valid events (*t*_btw_) (Fig. 2f), and a 1.5-fold decrease in the background emission frequency (*f*_bg_) (Fig. 2g).

In GLF-MINFLUX, each valid event encompasses labeling, localization, and bleaching, representing an individual target. We further employed GLF to 3D MINFLUX imaging to achieve single molecule separation across three dimensions, thereby providing detailed spatial and quantitative insights into densely packed structures. Utilizing 3D GLF-MINFLUX imaging of mitochondrial import receptor subunit TOM20 homolog, we revealed a laterally enriched TOM20 within the outer membrane of mitochondria (Fig. 2h,i, Extended Data Fig. 6, and Movie 3). Importantly, in each cluster, 3D GLF-MINFLUX identifies over 7.9 ± 6 HaloTag-fused TOM20 proteins, existing within spatial distances ranging from 2.7 to 18.5 nm in three-dimensional distribution (Fig. 2j,k).

**Fig. 2.**
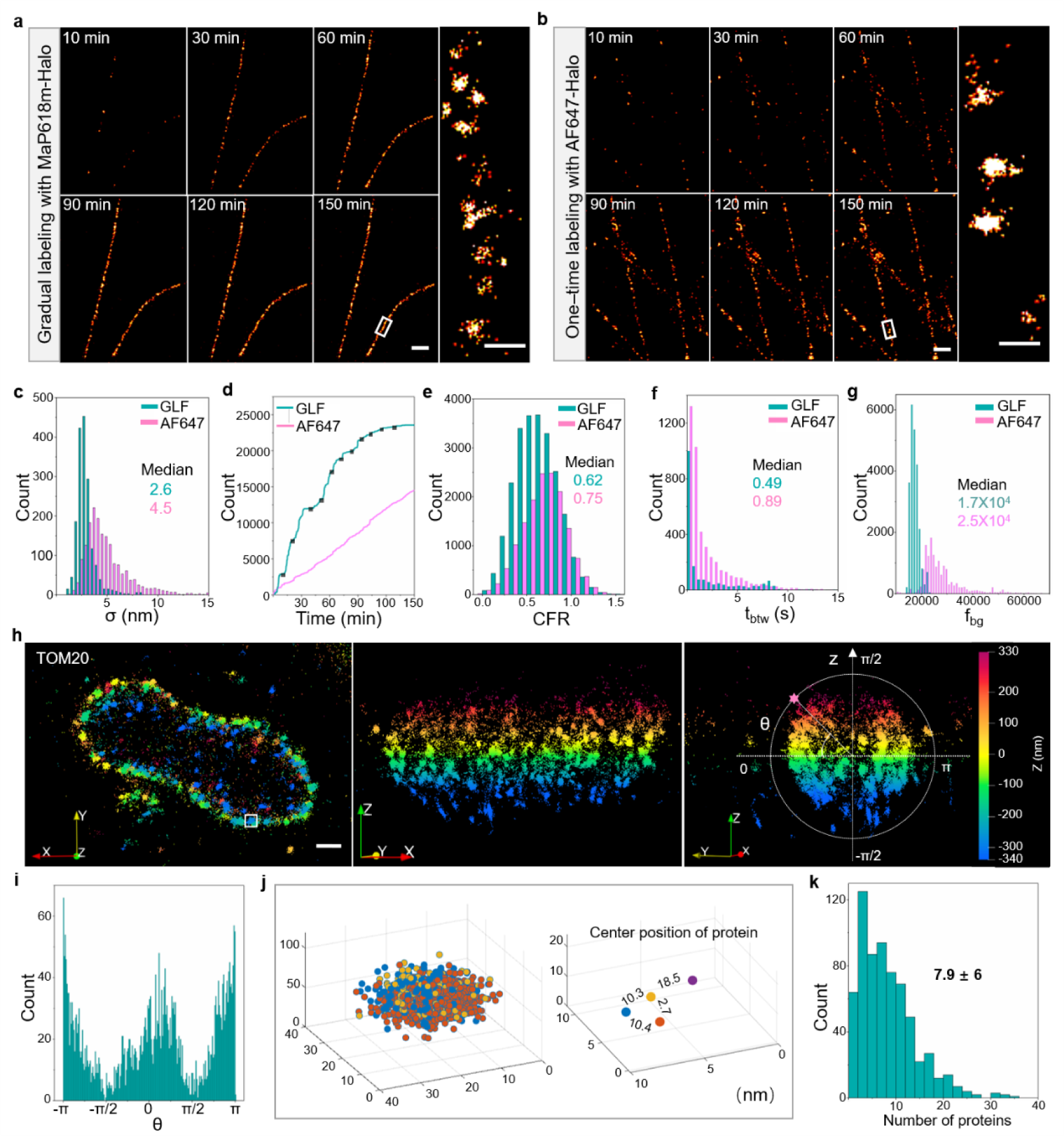
| D and 3D GLF-MINLUX nanoscopy of microtubules and mitochondrial TOM 0 clusters. **a**, GLF-MINFLUX imaging of microtubules in Halo-EGFP-tubulin expressing BS-C-1 cell. Individual panels show all recorded localizations in the indicated time intervals. **b**, MINFLUX imaging of microtubules after one-step labeling with Alexa Fluor 647 in Halo-EGFP-tubulin expressing BS-C-1 cell. The panel in the rightmost inset represents the zoomed region of microtubules within the white box in the panel 150-minute in **a** and **b**. **c**, Histograms of the localization precisions in microtubule images using GLF-MINFLUX and MINFLUX nanoscopy with Alexa Fluor 647 (N=5). **d**-**g**, The comparative analysis of GLF-MINFLUX and MINFLUX nanoscopy with Alexa Fluor 647 in terms of imaging speed (**d**), CFR (**e**), *t*_btw_(**f**), and *f*_bg_ (**g**). The black dots in **d** indicates the addition of fresh probes. **h**, 3D GLF-MINFLUX imaging of TOM20 in the mitochondria of HeLa cell. TOM20 fused with HaloTag was labeled with **MaP619m-Halo**. **i**, Angular statistics of TOM20 around the long axis of the mitochondria in **h**. **j**, The 3D distribution of the clusters of TOM20 within the white box in the plane **h**. The colored dots on the right represent the central points of the corresponding localizations on the left. **k**, Protein count of TOM20-HaloTag within each TOM cluster (N=8). Scale bars (MINFLUX) 500 nm (**a**, **c**), 200 nm (**h**). Scale bar (MINFLUX close-up), 50 nm (**a**, **c**).

Neuronal axonal cytoskeleton is vital for controlling cellular functions such as cell division, migration, intracellular trafficking, and signal transduction. However, the dense and intricately bundled arrangement of microtubules and microfilaments remains elusive in MINFLUX nanoscopy^14, 15^. Moreover, conventional large-size antibody-conjugated probes usually suffer from constrained imaging resolution and limited penetration into the confined interstitial spaces amidst bundled microtubules^16^. To address this issue, we conjugated ligands cabazitaxel and jasplakinolide with **MaP618m**, resulting in the development of small-molecule fluorogenic probes **MaP618m-tubulin** and **MaP618m-actin**, capable of directly binding to endogenous microtubules and microfilaments (Extended Data Fig. 1, Supplementary Table 2 and Note 2). These probes exhibit remarkable signal-to-noise ratios, typically ranging from 200 to 300-fold, upon binding to microtubules and microfilaments in both solution and fixed cells (Extended Data Fig. 7,8 and Supplementary Note 5). By using automated image processing software “Line Profiler”^17^, GLF-MINFLUX nanoscopy of microtubules labeled with **MaP618m-tubulin** in BS-C-1 cells revealed a peak-to-peak distance between microtubule sidewalls averaging 22.4 ± 7 nm (Fig. 3a), aligning well with the diameters in reported work^18^. Moreover, GLF-MINFLUX nanoscopy enables the observation of more than seven individual microtubule lines within a single neuron (Fig. 3b). Each microtubule line exhibits a diameter of 24 ± 5 nm, and the inter-line spacing measures 63 ± 17 nm ^19^. Additionally, the density heat map centered on each probe unveiled a tubulin density reaching up to 14,000/µm^2^ (Fig. 3c), significantly exceeding the theoretical maximum density of 6064/µm^2^ calculated through one-step labeling with Alexa Fluor 647 (Supplementary Note 1). Remarkably, the sequential gradual labeling of **MaP618m-tubulin** and **MaP618m-actin** enables the nanoscale imaging of microtubules and microfilaments in a dual-channel MINFLUX nanoscopy, employing just a single excitation laser (Fig. 3e,f).

**Fig. 3.**
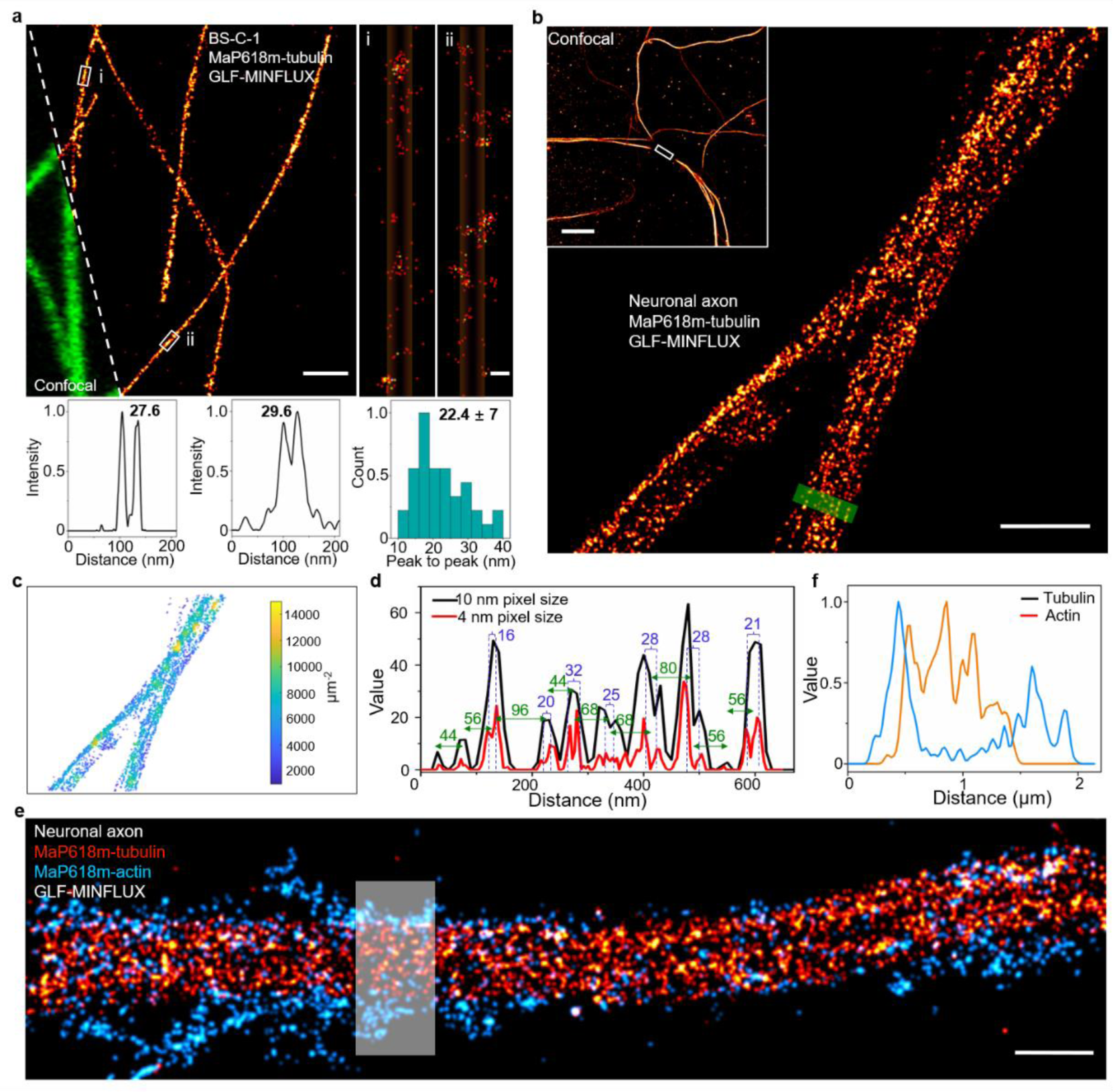
| GLF-MINLUX nanoscopy of densely packed neuronal cytoskeleton. **a**, GLF-MINFLUX imaging of microtubules in fixed BS-C-1 cell. The right panels depict zoomed-in imaging of microtubules within the white boxes on the left. Gaussian fitting curves of microtubules in panels i and ii and statistical analyses of peak-to-peak distances are presented at the bottom. **b**, GLF-MINFLUX imaging of microtubules in the axons of rat hippocampal neurons. The white outlined region in the top-left confocal image denotes the MINFLUX imaging field. **c**, Density heat map of the central positions of each emitter in **b**. **d**, Intensity profiles along the paths in the green box of **b**. **e**, Dual-channel MINFLUX nanoscopy of microtubules and microfilaments in the axons of rat hippocampal neurons labeled with **MaP618m-tubulin** and **MaP618m-actin**. Excitation laser wavelength: 642 nm. **f**, Intensity profiles of microtubules and microfilaments within the white boxed regions of **e**. Scale bars (MINFLUX) 1 µm (**a**, **b**, **e**). Scale bar (MINFLUX close-up) 20 nm (**a**). Scale bars (confocal) 10 µm (**b**).

In summary, GLF-MINFLUX offers a valuable avenue for nanoscale imaging of densely packed samples. This methodology effectively tunes the number of illuminated targets for MINFLUX nanoscopy by simply adjusting fluorogenic probe concentration. Compared to initial MINFLUX relying on the popular “on/off” switchable dyes Alexa Fluor 647, GLF-MINFLUX offers a 1.7-fold localization precision improvement and 2.2-fold faster acquisition while avoiding the use of complex buffer systems or dedicated activation laser. Crucially, GLF-MINFLUX allows for three-dimensional multi-target nanoscopy of densely packed structures through successive labeling with distinct probes even within the same color range. This innovative labeling method coupled with MINFLUX nanoscopy holds the promise of providing molecular-level insights into various biological samples.

## Methods

### Probe synthesis

Methods for organic synthesis and additional characterization can be found in the Supplementary Note.

### *In vitro* measurement of fluorescence spectroscopy

Absorption spectra and emission spectra were measured on BioTek Cytation 5 Cell Imaging Multimode Reader using a 384-well plate (Greiner, #781090) with optical bottom. **MaP618m-tubulin** (1 mM in DMSO) was diluted to 1 μM in the tubulin buffer or the presence of 2 mg/ml tubulin protein (cat. T240, Cytoskeleton Inc.) or 0.1% SDS. The tubulin buffer contained 80 mM PIPES pH 6.9, 2 mM MgCl2, 0.5 mM EGTA, 1 mM GTP (cat. BST06, Cytoskeleton Inc.), and 10% glycerol. The tubulin assay was incubated at 37 °C for 2 hours before absorption and emission spectra measurements were performed. The absorption spectra were collected from 400 to 700 nm with a slid width of 2 nm, while the emission spectra were obtained from 625 to 700 nm with a slid width of 2 nm under the excitation of 595 nm. All measurements were taken at ambient temperature (25 ± 2 °C) in general tubulin buffer unless otherwise noted.

### Plasmids and stable cell lines

Plasmid pLVX-EGFP-Halo-Tubulin and pLVX-TOM20-Halo-EGFP were stably transfected into BS-C-1 or HeLa cells via viral packaging, accompanied by two auxiliary plasmids, namely psPAX2 (plasmid #12260 Addgene) and pMD2.G (plasmid #12259 Addgene).

### Cell culture

BS-C-1 cells (ATCC, CCL-26, BeNa Culture Collection, China) were cultured in MEM (11095-080, Gibco) supplemented with 10% FBS (10099-141, Gibco), 100 U/mL penicillin, 100 µg/mL streptomycin (15140-122, Gibco), and non-essential amino acids (NEAA, 11140-050, Gibco). HEK293T (ATCC, CRL-3216, BeNa Culture Collection, China) and HeLa cells (ATCC, CCL-2, BeNa Culture Collection, China) were cultured in DMEM (11995-065, Gibco) supplemented with 10% FBS, 100 U/mL penicillin, and 100 µg/mL streptomycin. Cells were plated on cell culture dishes at 37°C with 5% CO_2_.

### Neuron culture

Hippocampal neuron cultures were derived from postnatal day P0–P3 Wistar rats of both sexes. The hippocampi were meticulously isolated and subjected to digestion with 0.25% trypsin-EDTA (25200072, Gibco) at 37℃ for 20 minutes. After centrifugation and removal of the supernatant, the cells were resuspended in the complete DMEM culture medium. Subsequently, the cells were plated on coverslips coated with 100 µg/mL of Poly-D-Lysine (A3890401, Gibco) according to the manufacturer’s recommendations. Neuronal cultures were maintained in Neurobasal-A medium (10888-022, Gibco) supplemented with 2% B27 serum-free supplement (17504044, Gibco), 1% Glutamax (35050-061, Gibco), and 100 U/mL penicillin and 100 µg/mL streptomycin. The neurons were fed with one-half medium volume change every 3 days.

### Cell fixation and labeling with fluorogenic probes

The cells were cultured for 1 day on 18 mm #1.0 coverslips (41001118, Deckglaser) positioned within 12-well cell culture dishes. Cells expressing HaloTag underwent a thorough rinsing with buffer, namely PBS for mitochondria imaging or extraction buffer (100 mM PIPES, 1 mM EGTA, 1 mM MgCl_2_, and 0.2% Triton X-100) for microtubule imaging. Following fixation with 2.4% paraformaldehyde (157-8, Electron Microscopy Sciences) and 0.1% glutaraldehyde (16220, Electron Microscopy Sciences) for 10 minutes, cellular entities experienced a triple PBS rinse, followed by a brief wash in PBS and immersion in a 10 mM sodium borohydride (71320, Sigma) solution in PBS for 5 minutes. Subsequently, the samples underwent a trifold PBS wash and the introduction of fiducial markers (gold nanorods, A12-40-980-CTAB-DIH-1-25, Nanopartz Inc.). The provided gold nanorod solution underwent a 1:3 dilution in PBS and sonication for 5–10 minutes. Coverslips were incubated with the nanorod solution and 1mM DL-dithiothreitol (DTT) for 15 minutes, with the removal of floating nanorods achieved through multiple PBS washes.

For **MaP618-tubulin** and **MaP618-actin** labeling, cells underwent permeabilization for 90 seconds utilizing the extraction buffer. Subsequently, cells were incubated with 0.2% glutaraldehyde in PEM (100 mM PIPES, 1 mM EGTA, and 2 mM MgCl_2_) for 15 minutes, followed by a brief wash in PEM and reduction in a 10 mM sodium borohydride in PEM for 5 minutes. Following this, the samples underwent three washes with PEM, and gold nanorods were introduced in PEM as previously described. Removal of floating nanorods was achieved through multiple washes with PEM. Before imaging, the coverslip containing the sample was affixed to the dish bottom with a central opening.

### MINFLUX imaging

HaloTag expressing cells that had been soaked in 1mM DTT in PBS or n PEM were immersed in the dish, securely affixing it to the specimen holder. Data acquisition took place on an Abberior MINFLUX microscope (Abberior Instruments) equipped with a 1.4 NA, 100× Oil objective lens, and avalanche photo-diodes. As mentioned earlier, calibration of the MINFLUX microscope was performed before utilization^1^. Utilizing Imspector Software (v.16.3.15636-m2205-win64-MINFLUX, Abberior Instruments), cells were identified and brought into focus using the 488 nm confocal scan of the microscope. Subsequently, the stabilization system of the microscope was activated, ensuring a stabilization precision typically below 1 nm. A region of interest was designated in the confocal scan image, and the 642 nm excitation laser power was set to 293 µW. The recording channel was adjusted within 650-685 nm. The background in this method is exceptionally low, with minimal dependence on pinhole diameter. Therefore, all data collection was performed using the default pinhole diameter of 0.78 AU. Subsequently, appropriate sequences were chosen for data acquisition. The fixed cells were stained with fluorogenic MaP dyes, with the concentration adjusted according to the field of view before imaging. Refreshing the dye solution every 10-15 minutes was essential to ensure imaging speed until the acquisition of new localizations nearly ceased. For fixed cells labeled with Alexa Fluor 647, sample preparation and MINFLUX imaging were carried out following the methodologies outlined in previous publications^20^.

### MINFLUX data analysis

Data exportation encompassed MSR, MAT, and NPY formats to enhance versatility in processing through diverse software platforms. The exported files comprised a comprehensive array of recorded parameters, encapsulating both valid localizations and discarded nonvalid attempts. MSR files facilitated the direct visualization of confocal outcomes and the presentation of MINFLUX localization distribution. MAT files were compatible with Matlab (2022b), enabling the automated computation of localization precision, CFR, *t*_btw_, *f*_bg_, and other parameters through customized analysis scripts. In all computational analyses, solely the data from the conclusive iteration of MINFLUX were utilized.

The experimental localization precision was estimated by aligning the mean value of all localization groups acquired from individual emission bursts. A Gaussian function was employed to fit the resulting histogram, providing an estimation of the localization precision (Fig. 2b). For reliability, only localization clouds containing a minimum of five localizations were employed to prevent underestimation of the localization error due to statistical bias. The characterization and specific calculation of the parameters CFR, *t*_btw_, and *f*_bg_ align with the methodologies previously delineated^4^.

### MINFLUX sequences

Data acquisition for the MINFLUX microscope adheres to a predefined set of parameters detailed in a text file (see **Imaging_2D_Nov2022.json and Imaging_3D_Nov2022.json** in the Supplementary data). These parameters delineate a sequential framework governing the iterative amplification of single-molecule events. The MINFLUX iteration process encompasses four iterations and one pre-localization iteration in two-dimensional space. In the three-dimensional domain, it entails nine iterations and one pre-localization iteration. Critical parameters for the 2D iteration sequence include:

| | $L$ (nm) | Minimal<br>photon count | CFR limit | Laser power<br>factor |
| --- | --- | --- | --- | --- |
| Prelocalization |  | 160 | Off | 1 |
| Iteration 1 | 288 | 150 | Off | 1 |
| Iteration 2 | 151 | 100 | 0.8 | 2 |
| Iteration 3 | 76 | 100 | 0.8 | 4 |
| Iteration 4 | 40 | 150 | Off | 6 |

Critical parameters for the 3D iteration sequence include:

| | $L$ (nm) | Minimal<br>photon count | CFR limit | Laser power<br>factor |
| --- | --- | --- | --- | --- |
| Prelocalization |  | 160 | Off | 1 |
| Iteration 1 | 288 | 400 | Off | 1 |
| Iteration 2 | 288 | 100 | Off | 1 |
| Iteration 3 | 288 | 50 | Off | 1 |
| Iteration 4 | 151 | 67 | 0.9 | 2 |
| Iteration 5 | 151 | 33 | Off | 2 |
| Iteration 6 | 76 | 67 | 0.8 | 4 |
| Iteration 7 | 76 | 33 | Off | 4 |
| Iteration 8 | 40 | 100 | Off | 6 |
| Iteration 9 | 40 | 50 | Off | 6 |

### Image rendering in two dimensions

In the context of sequential images, the utilization of Matlab scripts streamlines the aggregation of valid localizations across diverse time intervals, wherein the point count is converted into grayscale images featuring pixel dimensions of 10 nm (Fig. 2a,c) or 2 nm (Fig. 2a,c, close-up). The resultant images are subsequently processed and visualized using ImageJ. In the case of other images, valid localization events were delineated using Imspector Software and presented as 2D histograms with bin sizes of 10 nm (Fig. 3a,b,e) and 2 nm (Fig. 3a, close-up).

### Image rendering in three dimensions

Incorporating MINFLUX-derived data into ParaView (5.8.1-Windows-Python3.7-msvc2015-64bit), each localization was substituted with a Gaussian distribution and visually depicted with chromatic coding based on the z-axis as a parameter. Employing the 3D viewer within the software, images were systematically rotated to capture snapshots from diverse orientations (Fig. 3h), subsequently amalgamated into an animated format (Extended Data Movie 3). Consecutively, the MINFLUX dataset was transposed into Matlab, where localization clusters corresponding to distinct events were marked with unique colorations. The targeted regions of interest were then isolated and represented using Matlab’s 3D scatter plot, facilitating comprehensive exploration and analysis.

### Determining the central point of each emitter

Upon importing MINFLUX data into Matlab, a distinctive label (Tid) number was assigned to each event, facilitating the association of localizations bearing identical labels with the same event. Utilizing a script, the central point of each event’s localization cloud was automatically calculated, serving as an estimate for the central point of each emitter. Substituting the original localizations with these derived central points enabled the precise determination of the spatial positions and quantities of individual probes within the specimen (Fig. 2i–k and Fig. 3c).

### Quantifying mitochondrial protein distribution

Mitochondria were conceptualized as flat cylinders situated on the xy-plane. Employing customized software with coordinate adjustments (mito_pcd_editor.zip in the Supplementary data), the major axis of the cylinder was aligned with the x-axis. A parallel shift along the z-axis ensured that the circular center of the cross-sectional view coincided with the z-axis origin. Extraneous points along the mitochondrial circumference were systematically removed, resulting in a refined representation. The angles between each protein and the y-axis were computed based on the y and z coordinate information, where 0 or π indicated the two sides of the mitochondria, π/2 represented the upper surface, and -π/2 denoted the lower surface (Fig. 2i and Extended Data Fig. 3).

### Visualizing the density of the targets

The acquired central points of emitters were integrated into Matlab, and employing automated scripting, the distances from each emitter to its closest neighbor were computed. This process generated a density heatmap illustrating the concentration of targets per square micrometer (Fig. 3c).

## Acknowledgments

This investigation was supported by the National Natural Science Foundation of China (32171360 and 22107020 to L.W.), Zhongshan Hospital (2022ZSQD01 to L.W.), Basic Research Special Zone Plan of Shanghai (22TQ020), Science and Technology Commission of Shanghai Municipality (21DZ1100500), Shanghai Municipal Science and Technology Major Project, Shanghai Frontiers Science Center Program (2021-2025 No. 20), The National Key Research and Development Program of China (Grant No. 2021YFB2802000)

## Author contributions

M.G., L.Y., L.W., and J.W. conceived the original concept and initiated the work; L.Y. designed the experiments and performed imaging experiments; D.S. synthesized probes; L.Y., L.C., S.L, and J.G. analyzed the data; M.G., L.Y., L.W., J.W., J.M., and Q.Z. discussed the results; M.G., L.Y., L.W., and J.W. wrote the paper with input from all authors. All authors approved the manuscript.

---

**Extended Data Fig. 1.**
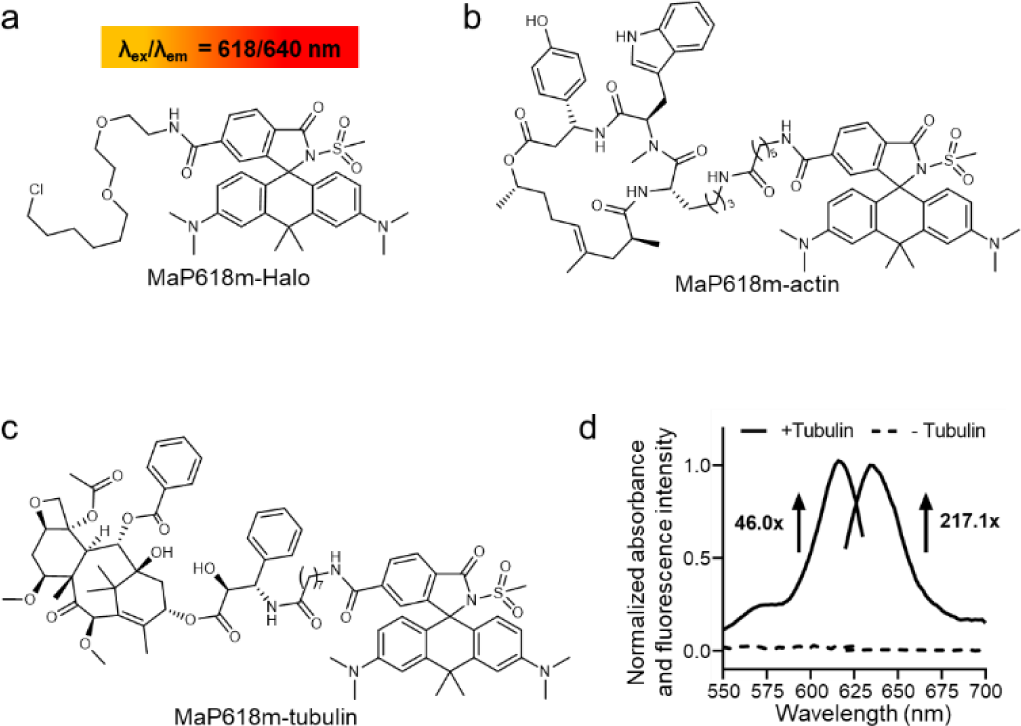
| Structures of fluorogenic MaP618m probes and the response of MaP618m-tubulin to microtubules. **a2** Structure of **MaP618m** coupled with chloroalkane for binding to HaloTag. **b**, Structure of **MaP618m** coupled with jasplakinolide for binding to F-actins. **c**, Structure of **MaP618m** coupled with cabazitaxel for binding to microtubules. **d**. Response of **MaP618m-tubulin** to microtubules. Absorption and emission spectra of **MaP618m-tubulin** (1 µM) were measured in the presence and absence of tubulin (2 mg/ml) after 2 h incubation at 37 ℃. The numbers in **d** indicate the ratio of absorbance at 616 nm and fluorescence intensities at 634 nm (λ_ex_ = 590 nm) in the presence and absence of microtubules.

**Extended Data Fig. 2.**
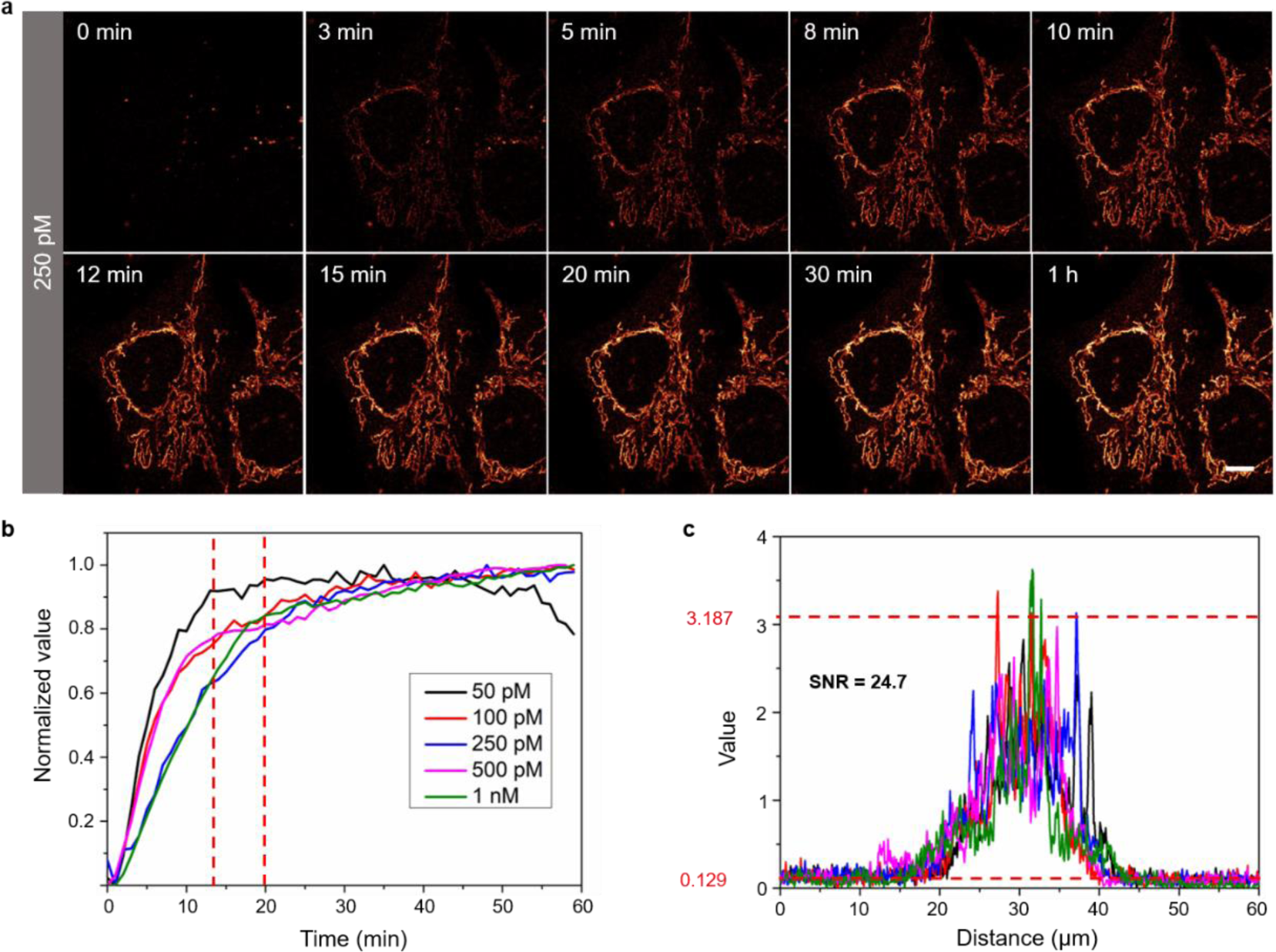
| Wash-free fluorescence images of BS-C-1 cells labeled with MaP618m-Halo. >**a**, Time-lapse confocal images of mitochondrial TOM20-HaloTag in fixed BS-C-1 cells labeled 250 pM **MaP618m-Halo** under wash-free conditions. Scale bar 10 µm. **b**, Fluorescent labeling of mitochondrial TOM20 in fixed BS-C-1 cells with **MaP618m-Halo** as a function of concentration. **c**, Singal-to-noise ratio (SNR) of mitochondrial TOM20 labeled with 250 pM **MaP618m-Halo** for 1 h. 5 cells were utilized for quantification. Excitation wavelength, 642 nm, detection range, 650 – 700 nm.

**Extended Data Fig. 3.**
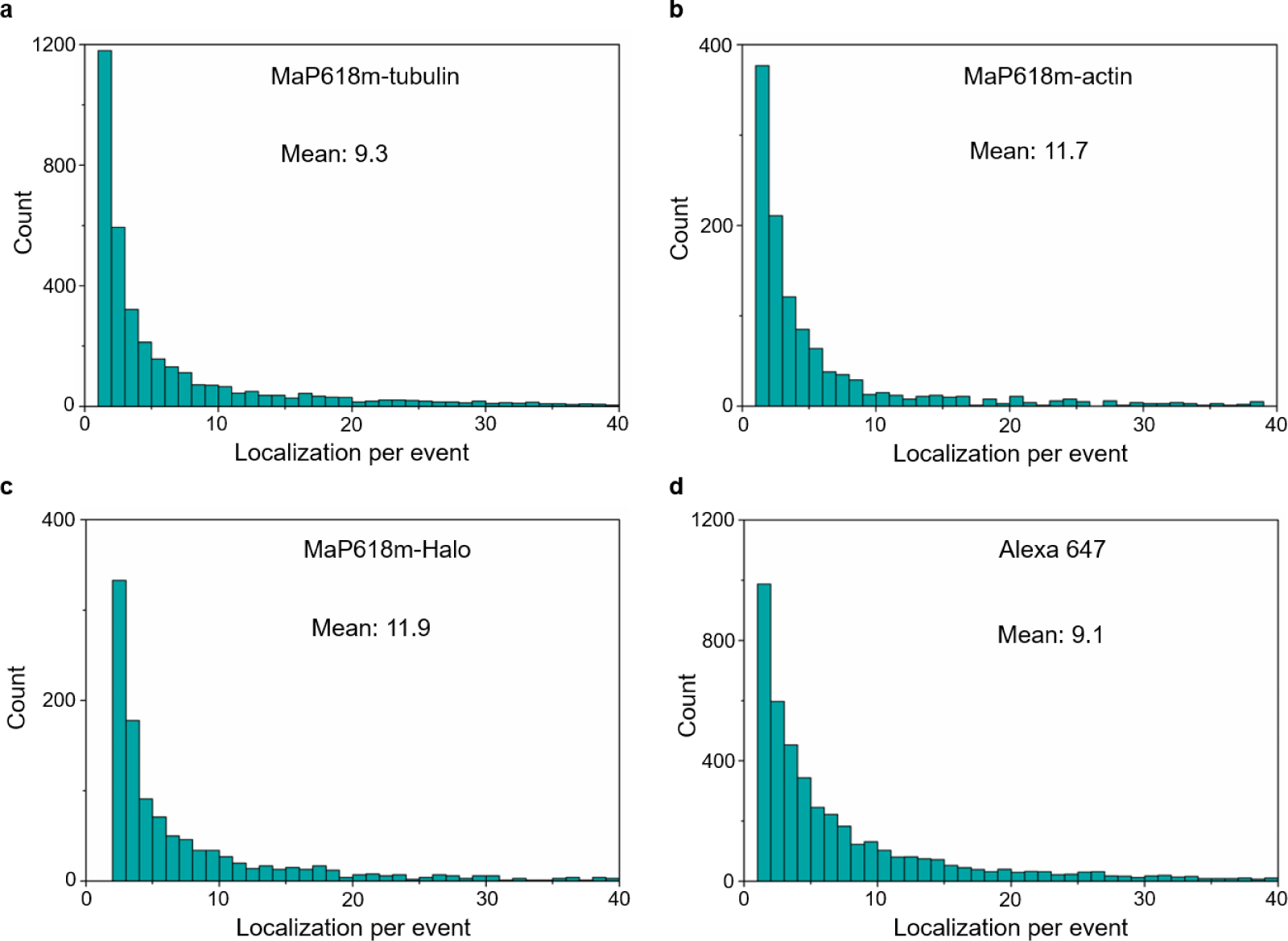
| The average number of localizations per localization cloud of MaP618m-tubulin2 MaP618m-actin2 MaP618m-Halo2 and Alexa Fluor 647. Halo-EGFP-tubulin expressing BS-C-1 cells were fixed and labeled with **MaP618m-tubulin**, **MaP618m-Halo2** or Alexa Fluor 647. And LifeAct-EGFP expressing BS-C-1 cells were fixed and labeled with **MaP618m-actin**. 10–16 μm^2^ ROIs were imaged for 2 hours each, using a laser power of 293 μW for **MaP618m** probes and 49 μW for Alexa Fluor 647 in the first iteration of MINFLUX imaging.

**Extended Data Fig. 4.**
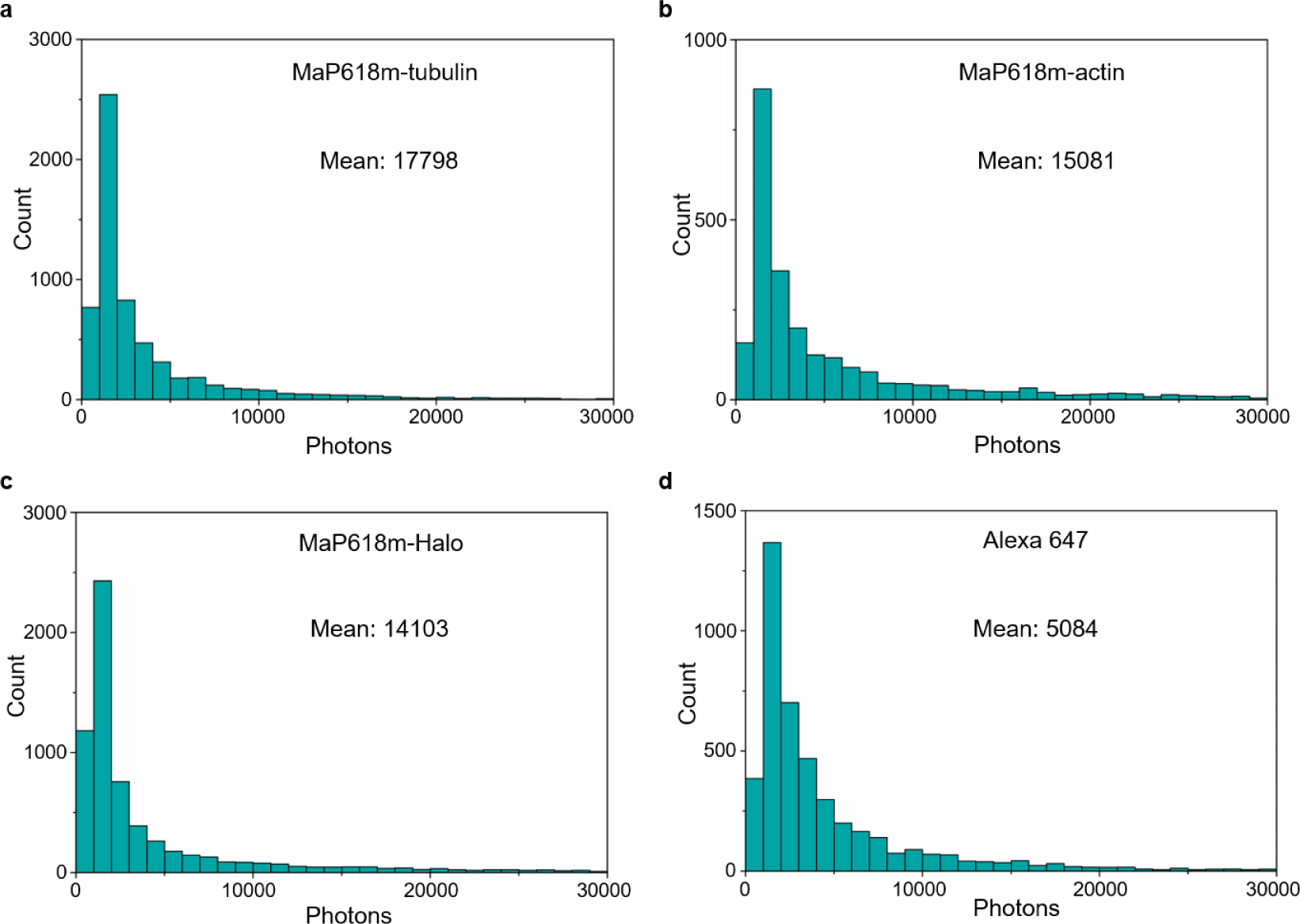
| Photon count statistics of MaP618m-tubulin2 MaP618m-actin2 MaP618m-Halo2 and Alexa Fluor 647. Halo-EGFP-tubulin expressing BS-C-1 cells were fixed and labeled with **MaP618m-tubulin**, **MaP618m-Halo**, or Alexa Fluor 647. And LifeAct-EGFP expressing BS-C-1 cells were fixed and labeled with **MaP618m-actin**. 10–16 μm^2^ ROIs were imaged for 2 hours each, using a laser power of 293 μW for **MaP618m** probes and 49 μW for Alexa Fluor 647 in the first iteration of MINFLUX imaging.

**Extended Data Fig. 5.**
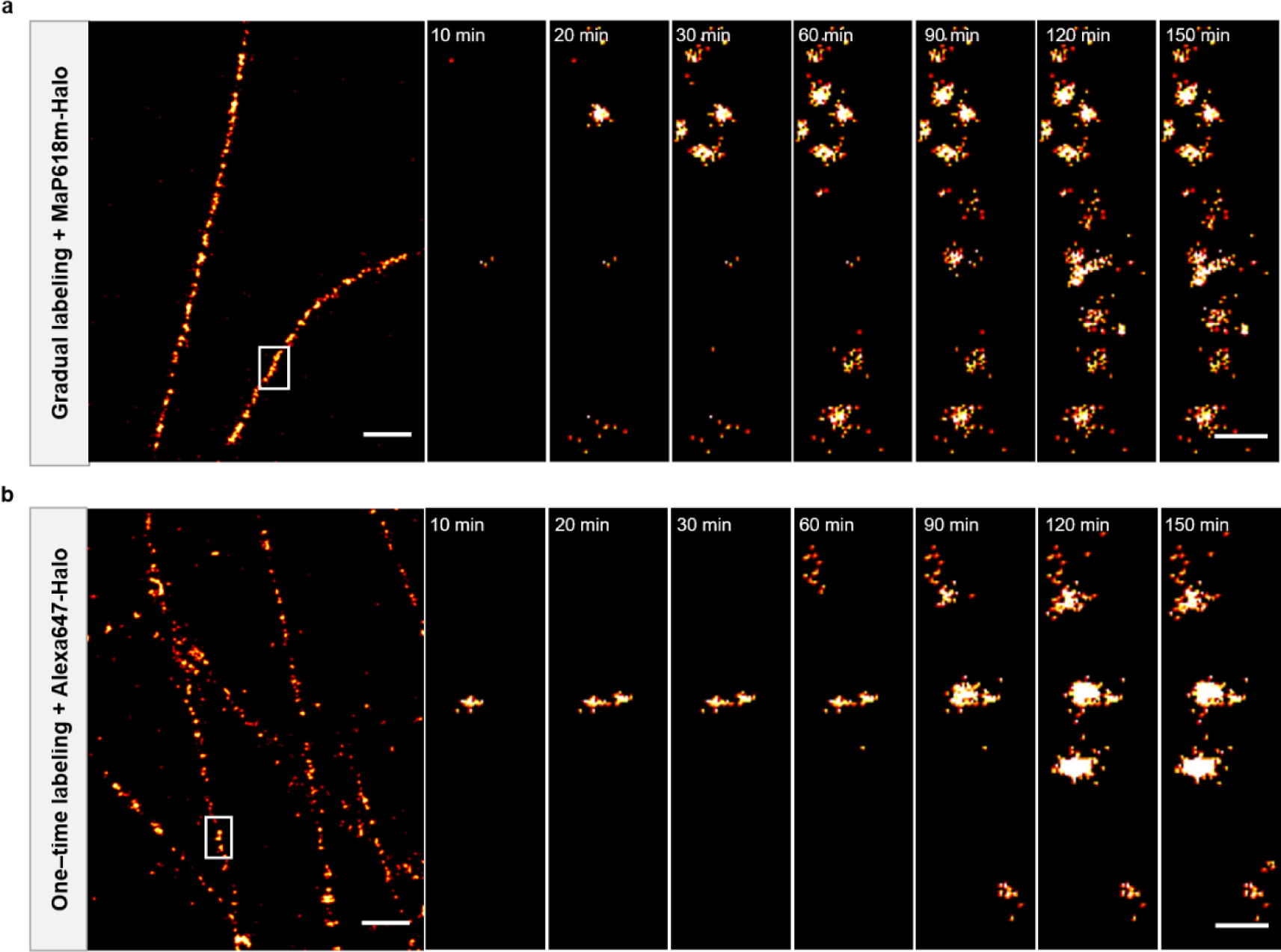
| MINFLUX nanoscopy of microtubules utilizing GLF-MINFLUX and Alexa Fluor 647. **a**, GLF-MINFLUX. **b**, Alexa Fluor 647-labeled MINFLUX. Halo-EGFP-tubulin expressing BS-C-1 cells were fixed and labeled with **MaP618m-Halo** or Alexa Fluor 647. 16.1 μm^2^ ROIs were imaged for 2.5 hours each, using a laser power of 293 μW for 100 pM **MaP618m-Halo** and 49 μW for Alexa Fluor 647 in the first iteration of MINFLUX imaging. Scale bars 500 nm (**a**, **b**). Scale bar (close-up) 50 nm.

**Extended Data Fig. 6.**
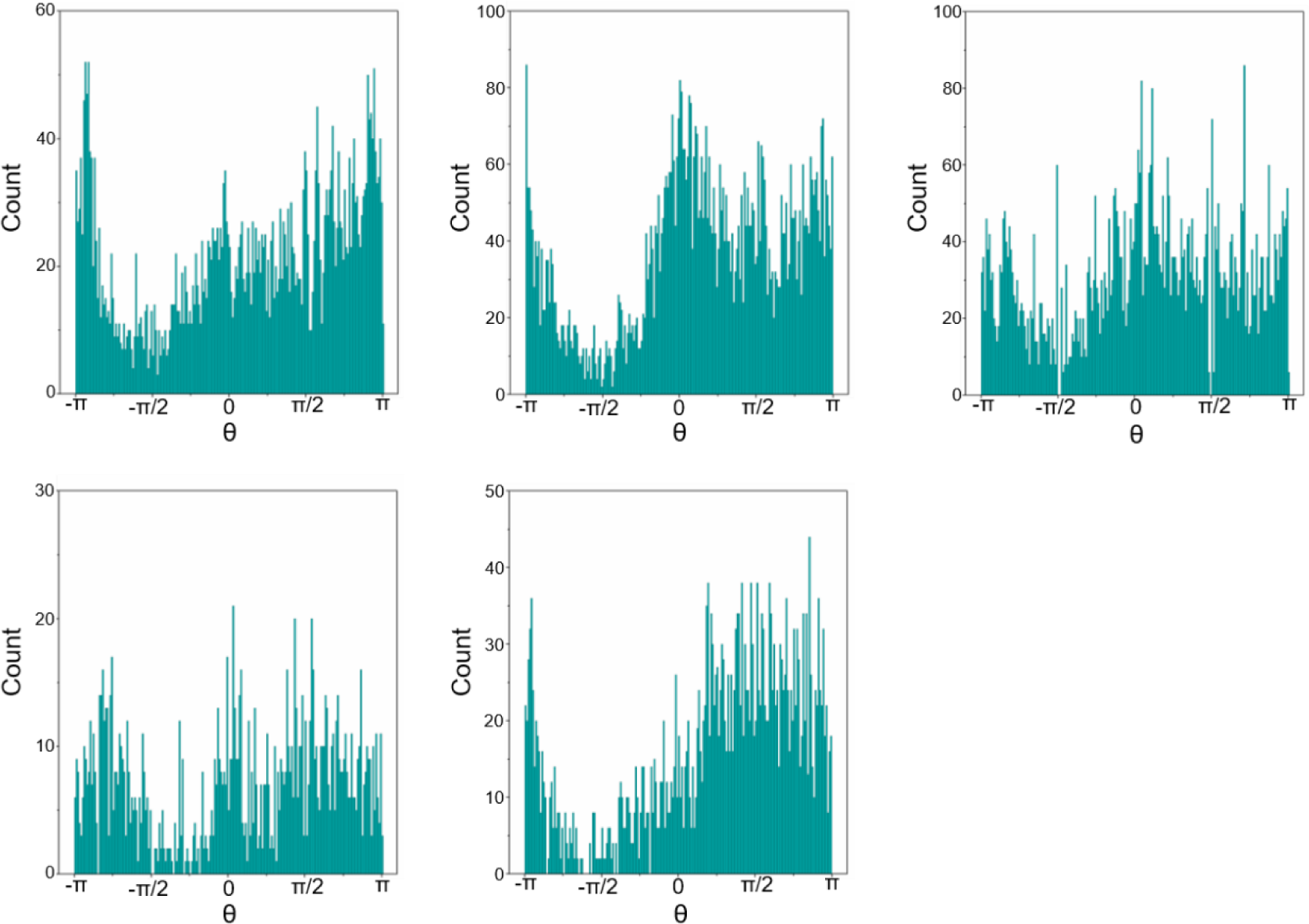
| The 3D spatial distribution of mitochondrial TOM 0 clusters. TOM20-Halo-EGFP expressing HeLa cells were fixed and labeled with **MaP618m-Halo**. 3–8 μm^2^ ROIs were imaged for 2–4 hours each, using a laser power of 293 μW for 100–150 pM **MaP618m-Halo** in the first iteration of MINFLUX imaging.

**Extended Data Fig. 7.**
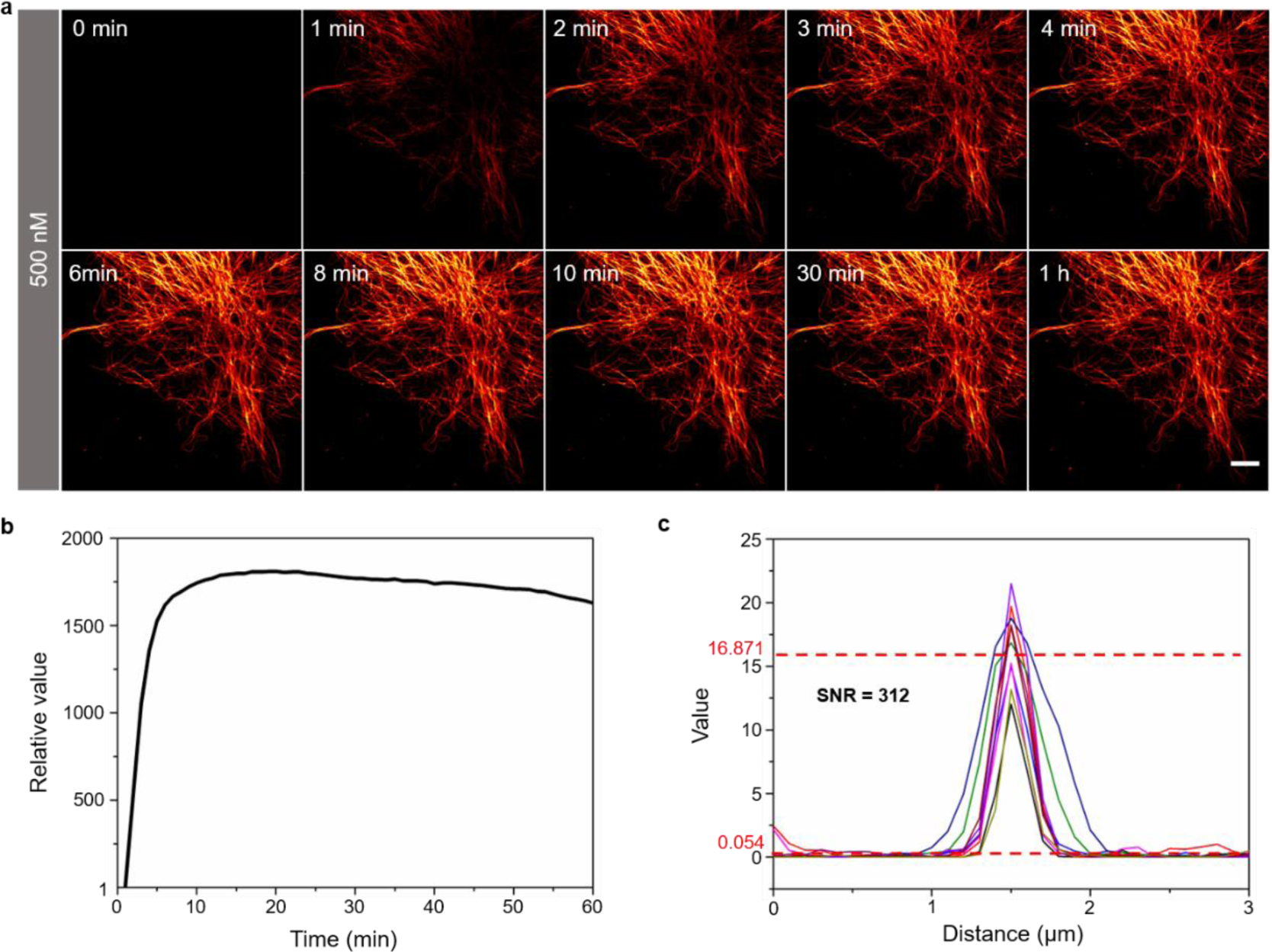
| Wash-free fluorescence images of BS-C-1 cells labeled with MaP618m-tubulin. **a**, Time-lapse confocal images of BS-C-1 cells labeled with 500 nM **MaP618m-tubulin** under wash-free conditions. Scale bar 10 µm. **b**, Labeling kinetic of microtubules in fixed BS-C-1 cells with 500 nM **MaP618m-tubulin**. **c2** Singal-to-noise ratio (SNR) of microtubules labeled with 500 nM **MaP618m-tubulin** for 1 h. 8 microfilaments in 4 cells were utilized for quantification. Excitation wavelength, 642 nm, detection range, 650 – 700 nm.

**Extended Data Fig. 8.**
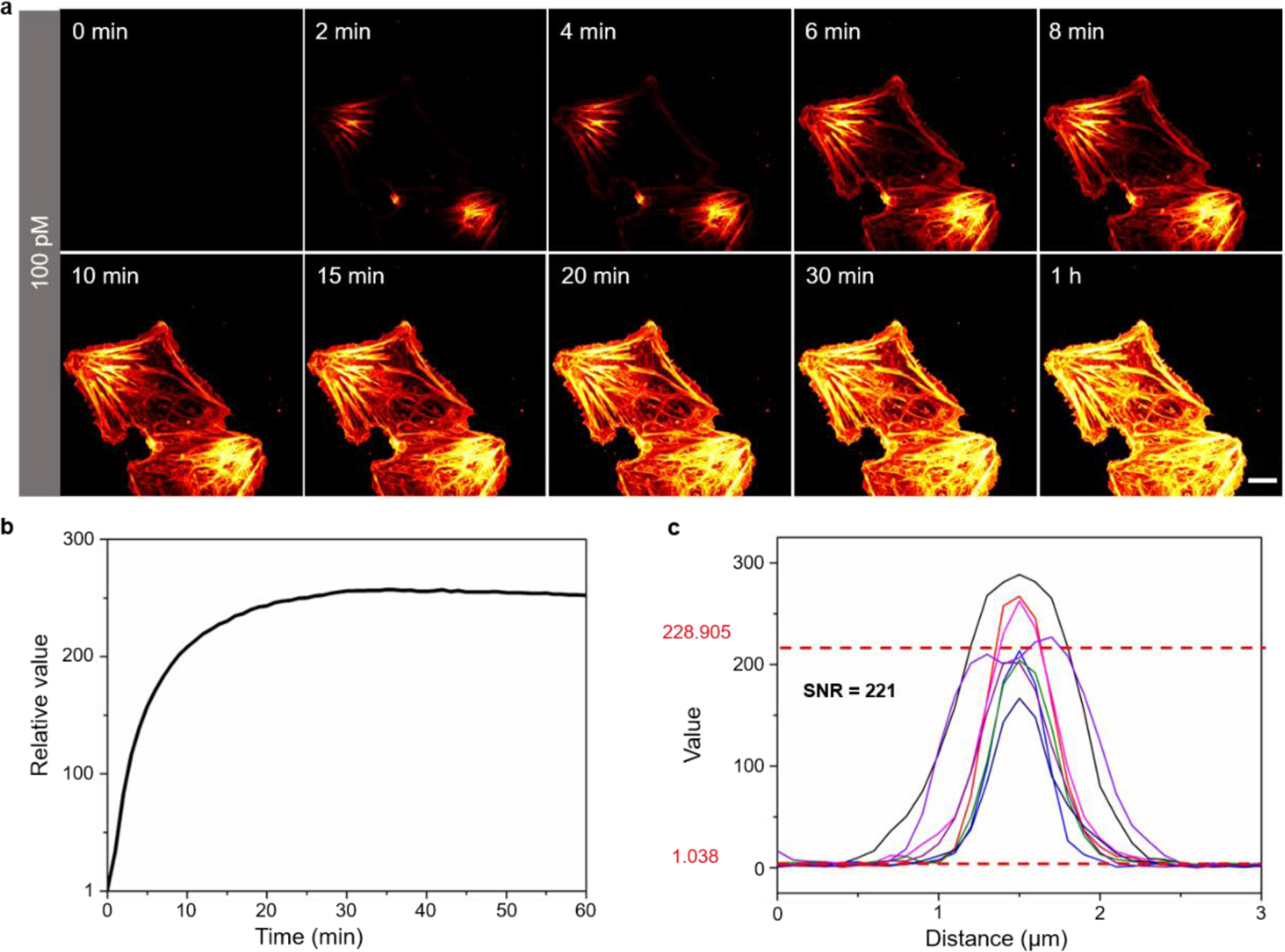
| Wash-free fluorescence images of BS-C-1 cells labeled with MaP618m-actin. **a**, Time-lapse confocal images of BS-C-1 cells labeled with 100 nM **MaP618m-actin** under wash-free conditions. Scale bar 10 µm. **b**, Labeling kinetic of microtubules in fixed BS-C-1 cells with 100 nM **MaP618m-actin**. **c2** Singal-to-noise ratio (SNR) of microtubules labeled with 100 nM **MaP618m-actin** for 1 h. 10 microtubules in 4 cells were utilized for quantification. Excitation wavelength, 642 nm, detection range, 650 – 700 nm.

